# Evasive spike variants elucidate the preservation of T cell immune response to the SARS-CoV-2 omicron variant

**DOI:** 10.1101/2022.11.04.515139

**Authors:** Arnav Solanki, James Cornette, Julia Udell, George Vasmatzis, Marc Riedel

## Abstract

The Omicron variants boast the highest infectivity rates among all SARS-CoV-2 variants. Despite their lower disease severity, they can reinfect COVID-19 patients and infect vaccinated individuals as well. The high number of mutations in these variants render them resistant to antibodies that otherwise neutralize the spike protein of the original SARS-CoV-2 spike protein. Recent research has shown that despite its strong immune evasion, Omicron still induces strong T Cell responses similar to the original variant. This work investigates the molecular basis for this observation using the neural network tools NetMHCpan-4.1 and NetMHCiipan-4.0. The antigens presented through the MHC Class I and Class II pathways from all the notable SARS-CoV-2 variants were compared across numerous high frequency HLAs. All variants were observed to have equivalent T cell antigenicity. A novel positive control system was engineered in the form of spike variants that did evade T Cell responses, unlike Omicron. These evasive spike proteins were used to statistically confirm that the Omicron variants did not exhibit lower antigenicity in the MHC pathways. These results suggest that T Cell immunity mounts a strong defense against COVID-19 which is difficult for SARS-CoV-2 to overcome through mere evolution.

**Author summary:** 

## 1 Introduction

The Omicron variant of SARS-CoV-2 caused the largest wave of COVID-19 infections from November 2021 to February 2022 [1]. Since then, Omicron has been responsible for more than 95% of all recorded cases of COVID-19 according to GISAID [2]. It was observed to have a higher infectivity rate and a lower disease severity than the original COVID-19 variant [3]. Notably, Omicron could reinfect patients who had already recovered from COVID-19 and infect people who had received double-doses of COVID-19 vaccines [3]. These factors had selected for its spread over other SARS-CoV-2 variants.

COVID-19 vaccines allow treated people to develop immunity against COVID-19 [4, 5]. The vaccines developed by BioNTech-Pfizer and Moderna-NIAID employ new mRNA vaccine technology, while those developed by Janseen-Johnson & Johnson, and Novavax feature previously established methods such as viral vectors and protein adjuvants. All of these vaccines utilize the novel Spike Glycoprotein of SARS-CoV-2 to activate a treated person’s immune system [6, 7]. They train the immune system to recognize the spike protein and develop antibodies to neutralize it. This means that upon COVID-19 infection, a vaccinated individual’s immune system can mount a stronger and faster defense. Vaccines also allow the body to produce T cells that kill COVID-19 infected cells through the Major Histocompatibility Complex (MHC) presentation pathways.

The MHC Class I and Class II pathways allow cells to present antigens derived from endogenous and exogenous proteins respectively [8]. These antigens are small peptides that are broken apart from the source protein and then strong bound by the MHC Class I or Class II proteins. Whenever a MHC protein binds a peptide, the resulting peptide-MHC complex is transported to the cell exterior such that the peptide can then be presented to other cells for immune surveillance. This antigen presentation allows for CD8^+^ and CD4^+^ T cells to identify any cells presenting pathogenic antigens and to consequently kill those infected cells. Clearly the MHC presentation pathways play a crucial role in immunity against diseases such as COVID-19.

MHC proteins of different alleles prefer binding different peptides [9]. For example, HLA-A*0201 prefers binding peptides with hydrophobic residues while HLA-B*2705 prefers binding peptides with hydrophilic residues. MHC Class I proteins also prefer binding peptides 9 amino acids long while Class II proteins prefer binding peptides of length 15. Furthermore, the MHC genes are among the most polymorphic genes in the human genome [10]. Due to all these reasons, the antigens presented from the SARS-CoV-2 spike protein by the MHC proteins can differ vastly from person to person. Individuals presenting a higher number of antigens elicit a stronger T Cell response to COVID-19 infection. Previous studies have concluded that the Omicron Variant is resistant to numerous monoclonal antibodies used in COVID-19 therapy, and evades antibodies from infected or vaccinated individuals as well [11–13]. However, T Cell responses to Omicron are mostly preserved in these individuals, suggesting that most T Cell epitopes from the original SARS-CoV-2 spike proteins are conserved in Omicron [14–19]. These observations motivate the importance of T Cell immunity in COVID-19 cases, as this class of immunity has not been strongly evaded by the SARS-CoV-2 virus with its new variants.

We sought to investigate the cause of this lack of evasion of T Cell Immunity by the Omicron variants. For our study, we used NetMHCpan-4.1 and NetMHCiipan-4.0 to predict MHC Class I and MHC Class II antigens respectively from various spike protein variants [20]. These are state-of-the-art tools that have been previously been used in vaccine design and neoantigen identification [21–23]. We used the predictions of these tools to investigate the antigens derived from the numerous SARS-CoV-2 Variants of Concern across various MHC alleles with high frequencies in human populations. These predictions estimated the different T-Cell responses elicited by the variants in most humans, and confirmed no significant loss of T-Cell epitopes in the Omicron spike protein. To validate this result, we generated a form of positive control by targeted the various epitopes we had identified in the spike protein, and deliberately mutated these regions of the protein to cause a loss of antigens binding to MHC proteins. These mutant “evading” spike proteins did lose T Cell epitopes across numerous HLAs when tested using NetMHCpan-4.1 and NetMHCiipan-4.0, unlike Omicron. These results prove that the Omicron spike protein lacks mutations that would lead to evasion of T Cell Immunity. This implies that other stronger selective factors (such as higher infectivity rate, stronger ACE2 binding, and antibody evasion) played a role in the evolution of the Omicron variant. Our results reassure that T Cell immunity forms a strong bastion against SARS-CoV-2, and should remain the focus of COVID-19 therapy.

## 2 Methods

### 2.1 SARS-CoV-2 Spike Protein and Variants

The spike protein in the original SARS-CoV-2 virus is a 1273 amino acid long protein [24, 25], and features a receptor binding domain which allows the virus to infect human cells through the angiotensin-converting enzyme 2 (ACE2) receptor [26]. Over the course of the COVID-19 pandemic from 2019 to 2022, the various Variants of Concern (VOCs) of SARS-Cov-2 reported by WHO evolved their own mutant spike proteins, with most mutations either stabilizing their spike protein or increasing its binding affinity with the ACE2 receptor. Clearly, the SARS-CoV-2 spike protein plays a key role in the COVID-19 viral infection, and consequently it has been the focus of numerous studies and clinical therapies. Notably, the most widely used COVID-19 vaccines in the United States (such as those manufactured by Pfizer, and Moderna) encode RNA transcripts for this spike protein, with minor stabilizing mutations. For our study, we sought to assess the various T-Cell antigens produced from spike protein variants from VOCs and common vaccines.

To begin, we identified all the key mutations with respect to the original spike protein in our different variants. For each of our SARS-CoV-2 variants, we recorded the deletions and point mutations that had a frequency of greater than 33% in all the spike protein samples for that variant in the GISAID database. We achieved this through the use of the mutation tracker on outbreak.info [27, 28] and [the list of shared mutations on the covariants website](https://covariants.org/shared-mutations) [29]. The mutations we observed across the different variants are listed below (listed in order of position along the spike protein):

1. B.1.1.7 (Alpha variant): H69del, V70del, Y144del, N501Y, A570D, D614G, P681H, T716I, S982A, and D1118H. There were 3 deletions and 7 point mutations.
2. B.1.351 (Beta variant): L18F, D80A, D215G, L241del, L242del, A243del, K417N, E484K, N501Y, D614G, and A701V. There were 3 deletions and 8 point mutations. Mutation L18F was not listed on the covariants website.
3. P.1 (Gamma variant): L18F, T20N, P26S, D138Y, R190S, K417T, E484K, N501Y, D614G, H655Y, T1027I, and V1176F. There were 0 deletions and 12 point mutations.
4. B.1.617.2 (Delta variant): T19R, T95I, G142D, E156G, F157del, R158del, L452R, T478K, D614G, P681R, and D950N. There were 2 deletions and 9 point mutations. Mutations T95I and G142D were not listed on the covariants website.
5. BA.1 (Omicron variant): A67V, H69del, V70del, T95I, G142D, V143del, Y144del, Y145del, N211I, L212del, G339D, S371L, S373P, S375F, K417N, N440K, G446S, S477N, T478K, E484A, Q493R, G496S, Q498R, N501Y, Y505H, T547K, D614G, H655Y, N679K, P681H, N764K, D796Y, N856K, Q954H, N969K, and L981F. There were 6 deletions and 30 point mutations.
6. BA.2 (“Stealth” Omicron variant): T19I, L24S, P25del, P26del, A27del, G142D, V213G, G339D, S371F, S373P, S375F, T376A, D405N, R408S, K417N, N440K, S477N, T478K, E484A, Q493R, Q498R, N501Y, Y505H, D614G, H655Y, N679K, P681H, N764K, D796Y, Q954H, and N969K. There were 3 deletions and 28 point mutations.

The mutations in the newest variants of concern of 2022, i.e. BA.2.75 and BA.5, are listed in the supplementary data 3.2.3. The majority of the mutations reported in these Omicron subvariants are also shared by BA.2.

We also tracked the synthetic mutations used in the common COVID-19 vaccines used in the United States. The vaccines we investigated were:

1. BNT162b2 (Manufactured by BioNTech-Pfizer) and mRNA-1273 (Manufactured by Moderna-NIAID): These feature the mutations K986P and V987P.
2. Ad26.COV2.S (Manufactured by Janssen-Johnson & Johnson): This features the mutations R682S, R685G, K986P, and V987P.
3. NVX-CoV2373 (Manufactured by Novavax): This features the mutations R682Q, R683Q, R685Q, K986P, and V987P.

With these key mutations compiled, we generated a consensus spike protein sequence for each variant by performing the corresponding single amino acid deletions or substitutions on the original spike protein. This statistics-based approach allowed us to investigate the impact of the core, defining mutations in a variant without relying on a single, incomplete sample to represent the variant. However this approach did not track amino acid insertions. For example, several BA.1 variant samples report small insertions in the N-Terminal Domain [30] but these could not be found in the mutation reporting data. Given this specific observation, we inserted the 3 amino acids EPE at position 214 in our BA.1 representative spike protein. This was the only insertion we performed in our analysis.

### 2.2 MHC-Peptide Binding Affinity Prediction

We used NetMHCpan-4.1 and NetMHCiipan-4.0 to investigate MHC Class I and Class II antigens respectively. Both of these tools predict the binding probability of a peptide with a given MHC molecule on a continuous scale of 0 to 1 – 0 means no affinity, and 1 means the strongest binding possible. Both these tools classify strong binding peptides to a MHC molecule using a 0.5 percentile threshold with respect to the training data for that MHC. For our MHC Class I analysis, we shortlisted the following HLA serotypes: A1, A2, A3, A24, A26, A30, B15, B35, B40, B44, and B51. For our MHC Class II predictions, we focused on DR1, DR3, DR4, DR7, DR8, DR11, DR12, DR13, DR1302, and DR15. We chose these HLA supertypes in our analysis because they are amongst the most frequent alleles while also representing a broad and diverse sample of the human population’s HLA types in the United States [31].

For each of our spike protein variants, we put the fasta-file sequence of the protein through both prediction tools, and gathered binding data for all the aforementioned MHC molecules. For MHC Class I predictions, we only analyzed 9mers – contiguous sequences of 9 amino acids generated from the protein – as they are the strongest binders for MHC Class I molecules compared to all other peptide lengths. For the same reason, we also focused on 15mers for MHC Class II molecules. The binding scores gathered from this prediction data allowed us to identify the antigens presented by various MHC molecules from each spike protein.

### 2.3 Generating Evaders as Positive Controls

In a peptide bound to a MHC protein, the starting and ending amino acids on the chain are tucked into the MHC’s binding pockets. The remaining peptide chain rests along a groove exposed on MHC surface, or bulges out if it is longer than can fit in the groove. Clearly, not every residue on the peptide contributes to its binding affinity with the MHC molecule. These certain positions on a strong binding peptide that play a crucial role in binding affinity are called *anchor residues*. When observing the binding motif of a particular HLA, i.e. the consensus sequence of all its strong binding peptides, anchor residues exhibit higher specificity for certain amino acids than the remaining positions do. Furthermore, different HLA types have different anchor residue preferences, due to biochemical factors such hydrophobicity, electrostatic interactions, and shape conformity that play a role in each HLA’s unique binding pocket. For example, A2 prefers binding 9mers with hydrophobic residues (L,V,M, and I) on positions 2 and 9, while B27 prefers binding 9mers with a simple R on position 2.

As highlighted in section 2.2, we needed to utilize some scientific controls to assess the significance of the number of predicted antigens being consistent across different variants. Our approach to resolve this was to design new spike protein variants as a form of positive control. These spike proteins would feature the same number of mutations as the most mutated SARS-CoV-2 variant. However, these spike proteins would be specifically engineered to knock down the number of antigens across multiple HLA types. Such spike proteins would act as positive controls since they would reflect the strongest MHC pathway knock down relative to all our tested variants. We termed such engineered spike proteins as “evaders”. We designed two sets of evaders – one for the MHC Class I pathway, and the other for Class II.

To design a Class I evader spike protein, we first identified which sites of the original spike protein to mutate to effectively knock down antigens. We mapped the location of all strong binding peptides from the original spike protein for each MHC Class I HLA along the 1273 amino acid sequence. This allowed us to identify *regions of antigenicity* – positions from the original spike protein that constitute peptides that are presented as antigens in the MHC Class I pathway. From here, we singled out the anchor residues in these regions of antigenicity. We compiled this information into a set of 11 lists – each list corresponding to an HLA type, and the numbers in each list representing the position of anchor residues for that HLA in the original spike protein. To generate the best evader protein, we needed to mutate these anchor residues. We limited the number of mutations to 36 (30 point mutations and 6 deletions), i.e. the number of mutations of BA.1. We also scaled the number of allotted mutations per HLA according to the number of antigens predicted for that HLA by NetMHCpan-4.1 (see Section 3.1.1). Therefore we randomly sampled 3, 3, 4, 3, 4, 4, 4, 5, 2, 2, and 2 numbers (a total sum of 36) from the 11 lists respectively. We then randomly mutated the 36 corresponding sites in the original spike protein to create an evader protein. For the point mutations, we ensured that the new amino acid substituting each chosen mutation site was antagonistic to the HLA that was anchored by that site. For example, a residue chosen for mutation that was originally an anchor residue for A2 binding was only allowed to mutate to amino acids besides V, L, M, and I. With this procedure we generated a 100 Class I evader proteins.

To design a Class II evader spike protein, we followed the same procedure to identify anchor residues. In this case, we had 10 lists and we sampled 6, 2, 3, 5, 2, 2, 3, 2, 6, and 5 numbers (again adding up to 36) from each respectively. We then repeated the same mutation protocol to engineer a 100 Class II evaders.

We repeated the procedure discussed in section 2.2 on the designed evaders to investigate the antigens predicted from them. We only looked at NetMHCpan-4.1 predictions for the Class I evaders, and NetMHCiipan-4.0 predictions for the Class II evaders.

### 2.4 Statistics and Ranking

For each of the 100 Class I evader proteins, we tracked the number of predicted strong binders by NetMHCpan-4.1 across all the 11 aforementioned HLAs. Here our goal was to analyze whether the 10 *nonrandom* variants (i.e. the original spike, the natural variants, and the vaccine variants) exhibited a different antigen profile compared to the evader proteins. To achieve this, we first used the Wilcoxon rank-sum test from the SciPy library to analyze whether any two sets of spike proteins possessed the same distribution of antigens. For each of the 11 HLAs, we compared the number of nonrandom variants’ predicted binders with the evasive variants’ predicted binders. Furthermore, we tested the BA.1 Omicron variant in comparison to both the nonrandom set and the evasive set. With the Wilcoxon test, we aimed to highlight that the set of nonrandom variants reported a different distribution of number of antigens for an individual HLA than the evasive proteins. Simply put, we tested if the evasive variants reported a different number of antigens across the various HLAs.

However, this test only investigated the differences between the nonrandom and evasive variants while focusing on an individual HLA. To understand the significance across all the HLAs, we utilized 3 different multi-HLA metrics as well. The first metric was a simple sum – for each variant we added up all the antigens across different HLAs to track the *total number of antigens*. This metric was useful for visualizing how many antigens were being lost over different evaders (the larger the total, the more the antigens), but was mostly determined by the few HLAs that presented many more antigens than others. So, we used the *cartesian distance* between a variant and the original spike. We treated each variant as a data point with each HLA as a coordinate and computed the root of the sum of the squares of its difference from the original spike. The larger the cartesian distance, the more different the antigen profile of a variant from the original spike. This metric did not scale down different HLAs with greater antigen variance, so we also developed a third rank metric to merge the multidimensional HLA data. For each individual HLA, we ranked all the spike proteins (from the 110 total nonrandom and evasive variants) based on the number of predicted antigens for that HLA in ascending order. Proteins that shared the same number of binders were assigned the same average rank. With all 110 proteins being ranked across all 11 HLAs, we added up the 11 ranks for a single protein into a single *sum of ranks* score. We repeated the Wilcoxon rank-sum test on all 3 of these metrics to identify if the antigen profiles of the nonrandom and evasive variants were distinguishable when considering all HLAs. Again, we investigated which set Omicron BA.1 matched better with as well.

We repeated the analysis discussed above for the 100 Class II evader proteins across the 10 Class II HLAs as well.

## 3 Results

### 3.1 MHC Class I

#### 3.1.1 NetMHCpan-4.1 Results on Spike Protein Variants

For each of our constructed spike protein variants, the number of antigens as predicted by NetMHCpan-4.1 is shown in Fig. 1. The results for the vaccines and the newer Omicron subvariants are presented in Table 6. For each of the 11 Class I molecules, these numbers of antigens were mostly unchanged across the different spike variants (the largest drop was observed for A26 between the original spike protein and BA.1 – a loss of 3 binders from the original 22, which represents a 13% drop in number of antigens). That is, no particular variant exhibited a significant knockout of predicted Class I antigens compared to the original spike protein’s antigens. In particular, the two Omicron variants did not exhibit a drop in total number of antigens for molecules A1, A2, A3, A24, A30, and B44. In some cases (namely for A26, B35, and B51) the Alpha, Beta, and Gamma variants even gained more predicted antigens over the original spike protein. This suggests that the number of antigens derived from the different spike proteins for an individual Class I HLA’s predictions is relatively consistent. The mutations present in newer variants, including Omicron, did not lower the number of antigens presented by the Class I pathway.

**Fig 1.**
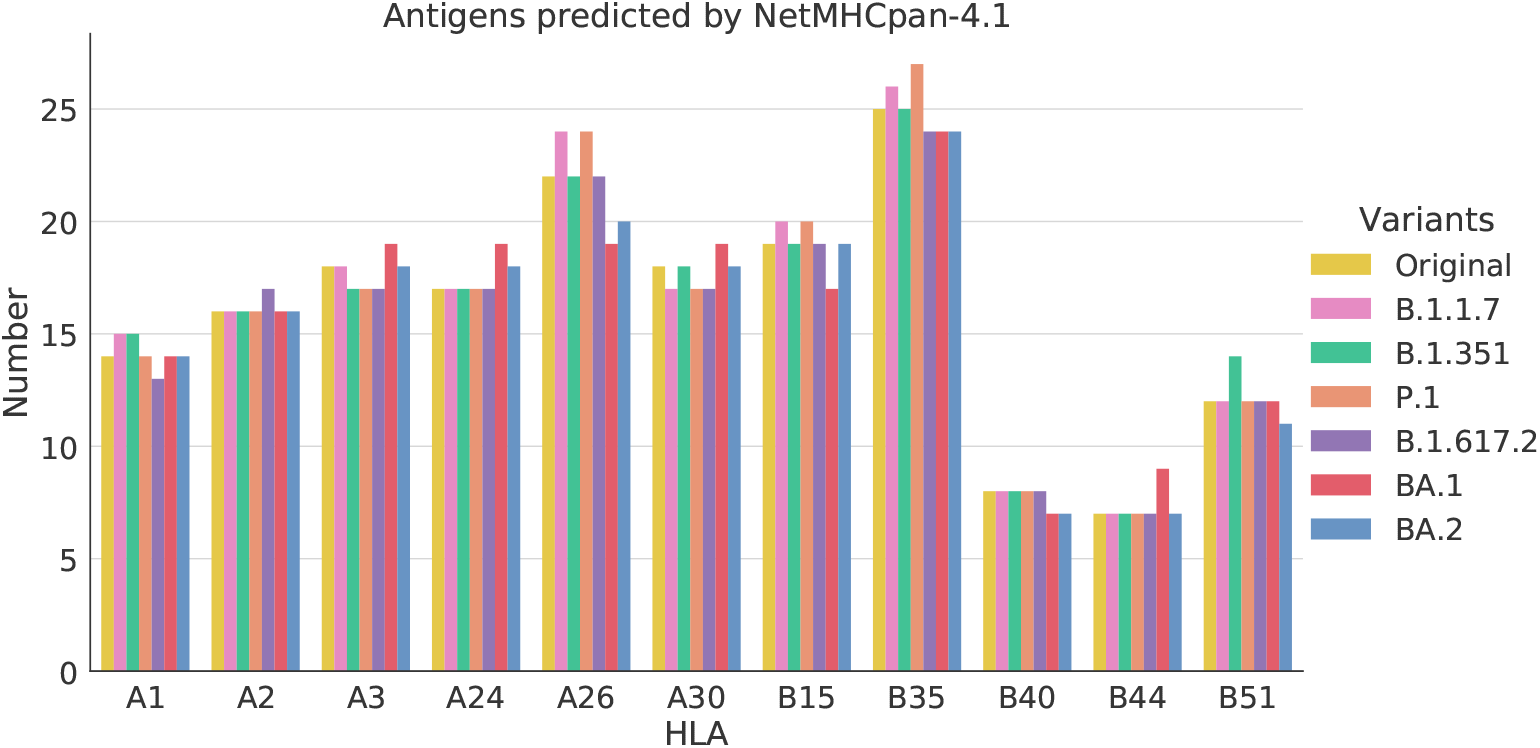
MHC Class I antigens predicted from the multiple SARS-CoV-2 variant proteins by NetMHCpan-4.1. The x-axis tracks the various common Class I HLA types while the y-axis reports the number of antigens. The results for different variants (from the original spike to both omicron variants) are shown in different colored bins for each HLA.

We also investigated the number of original SARS-CoV-2 peptides that were preserved in the newer variants. That is for each spike variant, we counted the number of its antigens that were also present in the original spike. These results are shown in Fig. 2 and Table 7. The two Omicron variants show a drop in the number of antigens across all HLAs. This means that certain mutations in these variants occur in regions of antigenicity of the spike protein. However, since the total number of antigens from these variants is equivalent to the original spike protein’s antigens (see Fig. 1), these mutations do not knockout the antigenicity of the spike protein. Instead, they merely alter the antigen footprint of the spike protein variants without actually impacting their net antigenicity.

**Fig 2.**
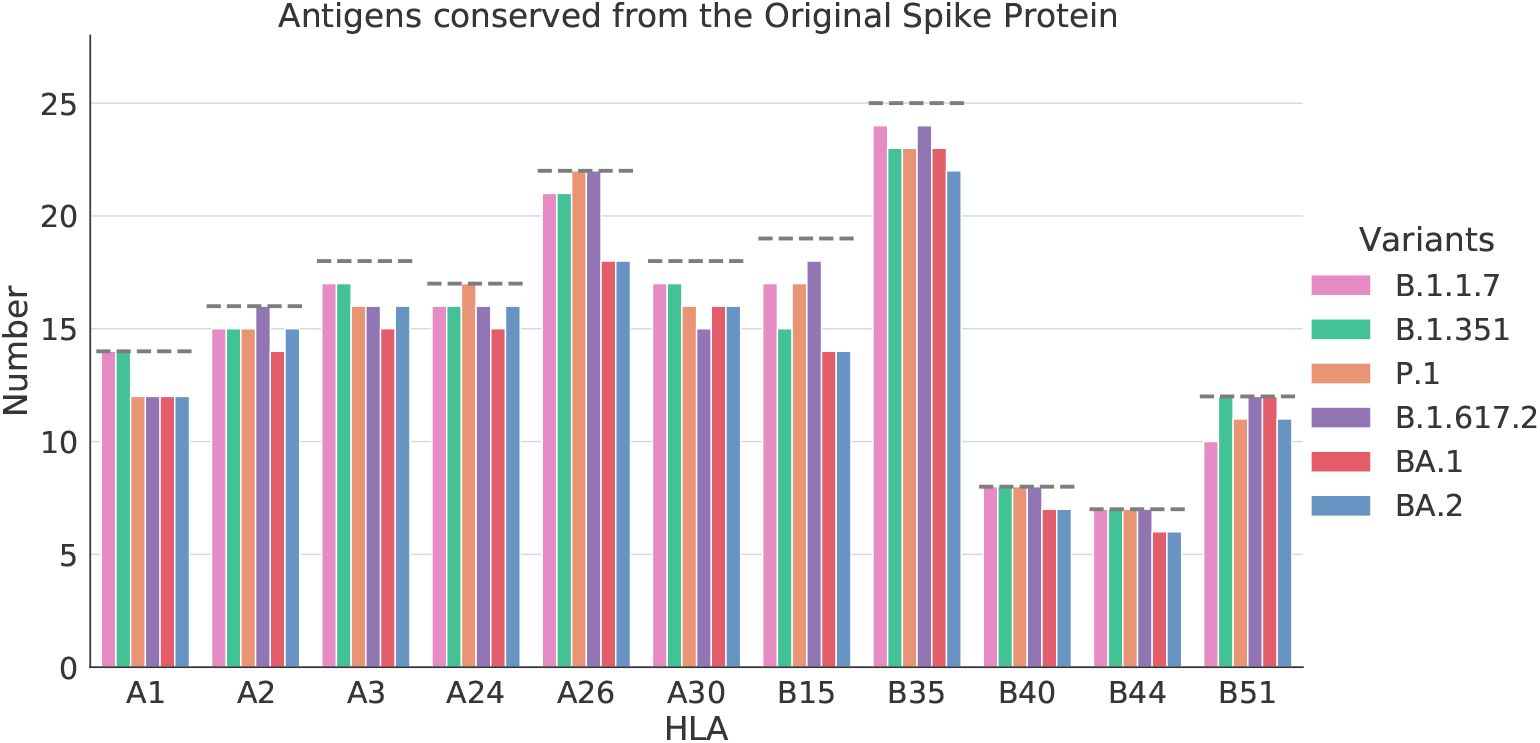
MHC Class I antigens from the original spike that were preserved in each variant spike according to NetMHCpan-4.1. For each HLA on the x-axis, the dashed line represents the number of antigens predicted from the original spike. The different variants are represented by the different colored bins. Each bin measures how many of the original spike antigens were conserved in a variant’s predicted set of antigens.

#### 3.1.2 NetMHCpan-4.1 Results on Evaders

The numbers of Class I antigens that were predicted by NetMHCpan-4.1 from our set of a 100 Class I evader proteins are shown in Fig. 3. For each HLA, the number of antigens counted in both the original spike and the omicron BA.1 spike are also reported. Furthermore, the mean numbers of antigens from the evaders are also shown. Clearly for all HLAs, the set of evaders reports a consistently lower number of antigens compared to both natural spike proteins. These results suggest that the targeted random mutations used to construct the evaders knocked down the antigenicity of the evader spikes across multiple HLAs. Note that since the 36 targeted mutations in an evader protein were distributed to lower antigenicity across all HLAs, no complete knockout of antigens is observed for a single HLA. Instead, a notable (but not complete) knockdown is observed for all HLAs. In summary, the mutations we sampled contributed to successful MHC Class I evasion by the evader proteins.

**Fig 3.**
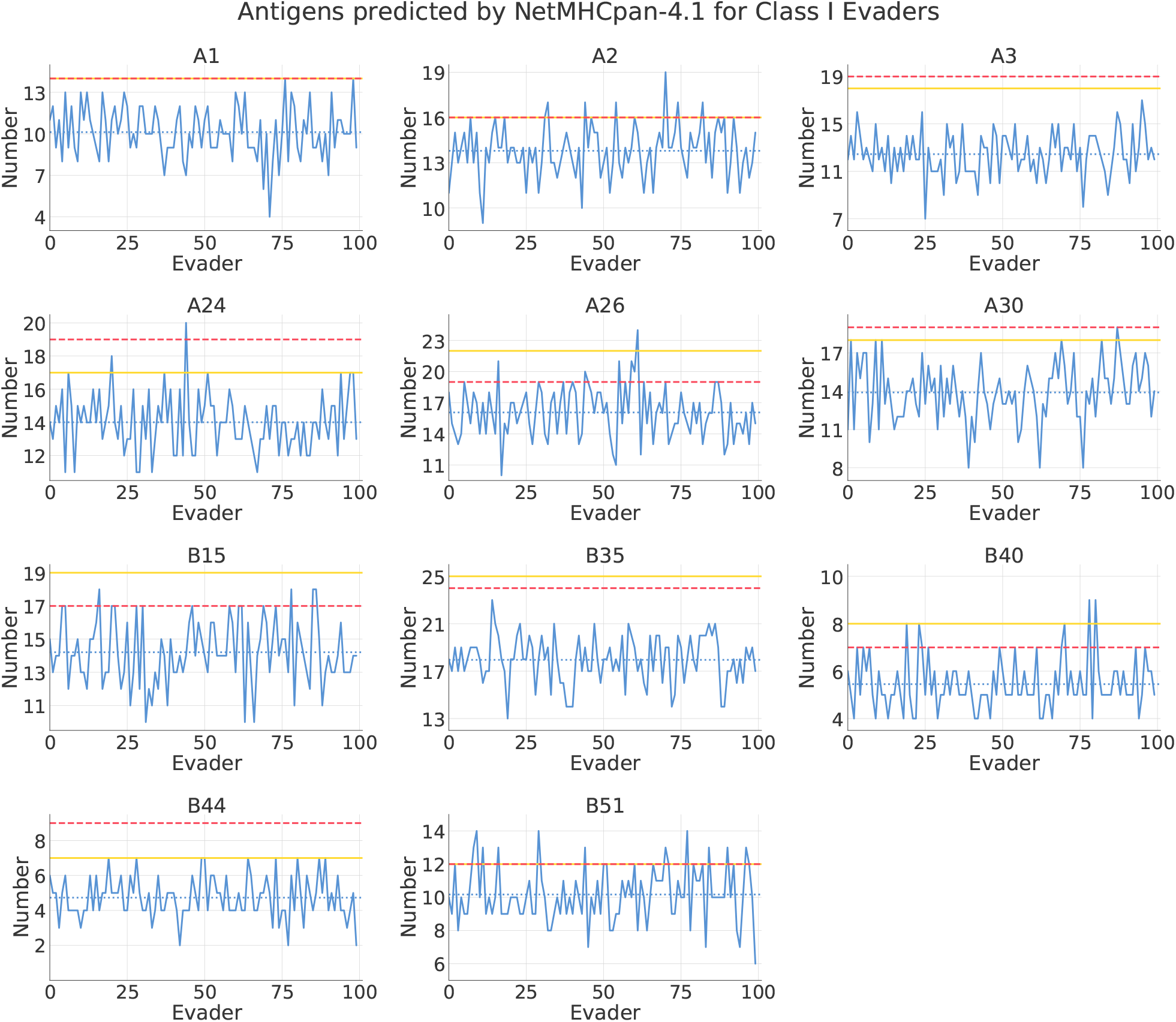
The number of strong binding peptides from each Class I evader protein predicted by NetMHCpan-4.1 for all target HLAs. The x-axis represents all 100 evaders and the y-axis represents the number of antigens. The evader predictions are shown in the blue line plot, with the mean number of antigens for all evaders in dotted blue. The number of predicted binders from the original spike is shown in solid yellow, and the number from Omicron BA.1 is shown in dashed red.

#### 3.1.3 Statistical Significance

We used the Wilcoxon rank-sum test to quantify how successfully our evaders lowered antigenicity of their spike proteins. The results for our single HLA tests are shown in Table 4. For all 11 HLAs, we confirmed that the set of evader proteins and the set of nonrandom proteins (natural variants and vaccines) formed statistically different distributions. This suggests that our evaders significantly knocked down antigenicity of the spike protein for each HLA. Furthermore, for each case the omicron variant compared more favorably with the nonrandom set than the evader set.

We also investigated 3 different multi-HLA metrics (as discussed in Section 2.4) to ensure our observations for single HLAs also applied for the entire catalogue of MHC Class I molecules. The distribution of these metrics for all evaders and nonrandom proteins is shown in Fig. 4. The first plot shows that the total antigens count, i.e. the sum of antigens from all tested HLAs, of all the evaders were lower than all nonrandom proteins. The second plot shows that all evaders had a higher cartesian distance from the original spike protein’s antigen profile in comparison to the other nonrandom variants. Lastly, the third plot depicts that all evaders possessed a lower sum of ranks score than the nonrandom proteins. All three of these metrics therefore confirm that the evaders knockdown antigenicity of the spike protein across all tested HLAs simultaneously. We utilized the Wilcoxon rank-sum test on these metrics and achieved the same conclusion as the single HLA tests, as shown in Table 1. That is, the Class I evaders bear mutations that consistently knock down the antigenicity of the spike protein across a whole catalogue of Class I MHCs. In fact, these evaders achieve these results while possessing the same number of mutations (36) as Omicron BA.1. Therefore the mutations seen in Omicron (and all other natural variants discussed in the paper) do not contribute to evasion of the MHC Class I pathway in one or many HLA alleles.

**Fig 4.**
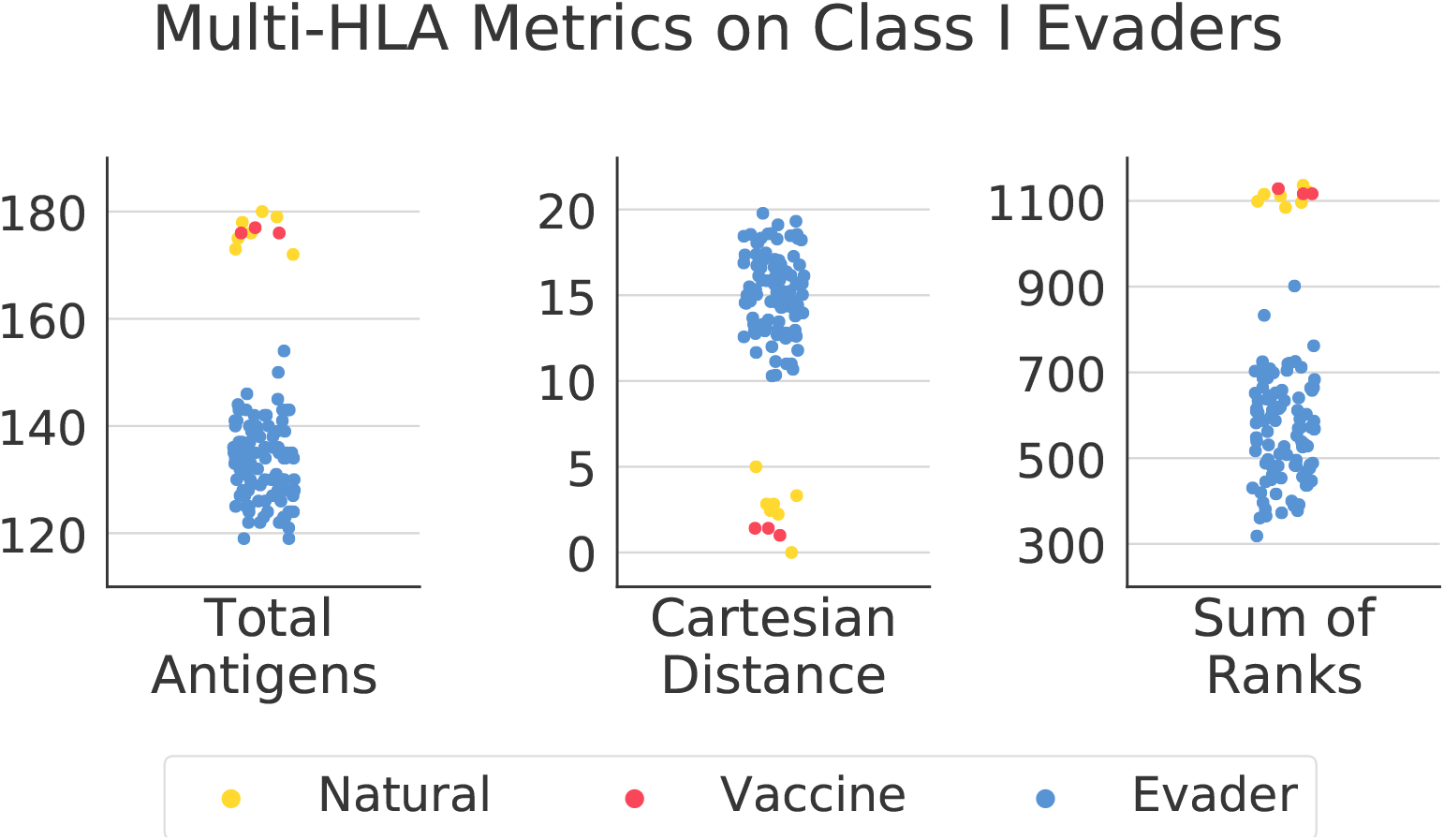
The distribution of the spike protein variants using the total antigens, cartesian distance, and sum of ranks metrics respectively. All Class I evaders are shown in blue, all natural SARS-CoV-2 are shown in yellow, and all vaccine spikes are shown in red. For each metric, the distributions of the evader proteins and the nonrandom (natural plus vaccine) are notably disparate.

**Table 1.**
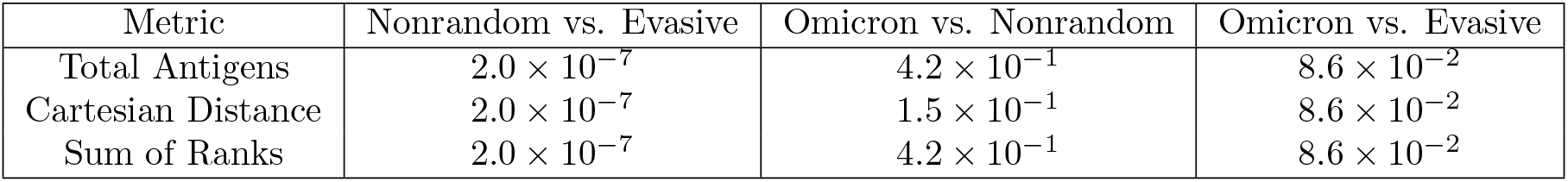
P-values for the Wilcoxon rank-sum test for multi-HLA metrics in Class I.

| Metric | Nonrandom vs. Evasive | Omicron vs. Nonrandom | Omicron vs. Evasive |
| --- | --- | --- | --- |
| Total Antigens | $2.0 \times 10^{-7}$ | $4.2 \times 10^{-1}$ | $8.6 \times 10^{-2}$ |
| Cartesian Distance | $2.0 \times 10^{-7}$ | $1.5 \times 10^{-1}$ | $8.6 \times 10^{-2}$ |
| Sum of Ranks | $2.0 \times 10^{-7}$ | $4.2 \times 10^{-1}$ | $8.6 \times 10^{-2}$ |

### 3.2 MHC Class II

#### 3.2.1 NetMHCiipan-4.0 Results on Spike Protein Variants

The prediction results of NetMHCiipan-4.0 for all our variants are shown in Fig. 5. Again, the number of predicted strong binders across different variants is relatively equivalent for each HLA allele (the largest drop was observed for DR4 between the original spike protein and B.1.1.7 – a loss of 5 binders from the original 19, which represents a 26% drop in number of antigens). For DR1, DR8, DR11, DR12, DR13, and DR1302, the two Omicron variants did not lead to a loss in number of antigens compared to the original spike protein. In the case of DR11, which bound the least number of antigens from the original spike at 2, all natural variants (i.e. Alpha through both Omicrons) did not yield fewer antigens. Thus similarly to the Class I predictions, the Class II predictions suggest that the mutations occurring in the natural SARS-CoV-2 variants did not lead to loss of presented antigens compared to the original spike protein.

**Fig 5.**
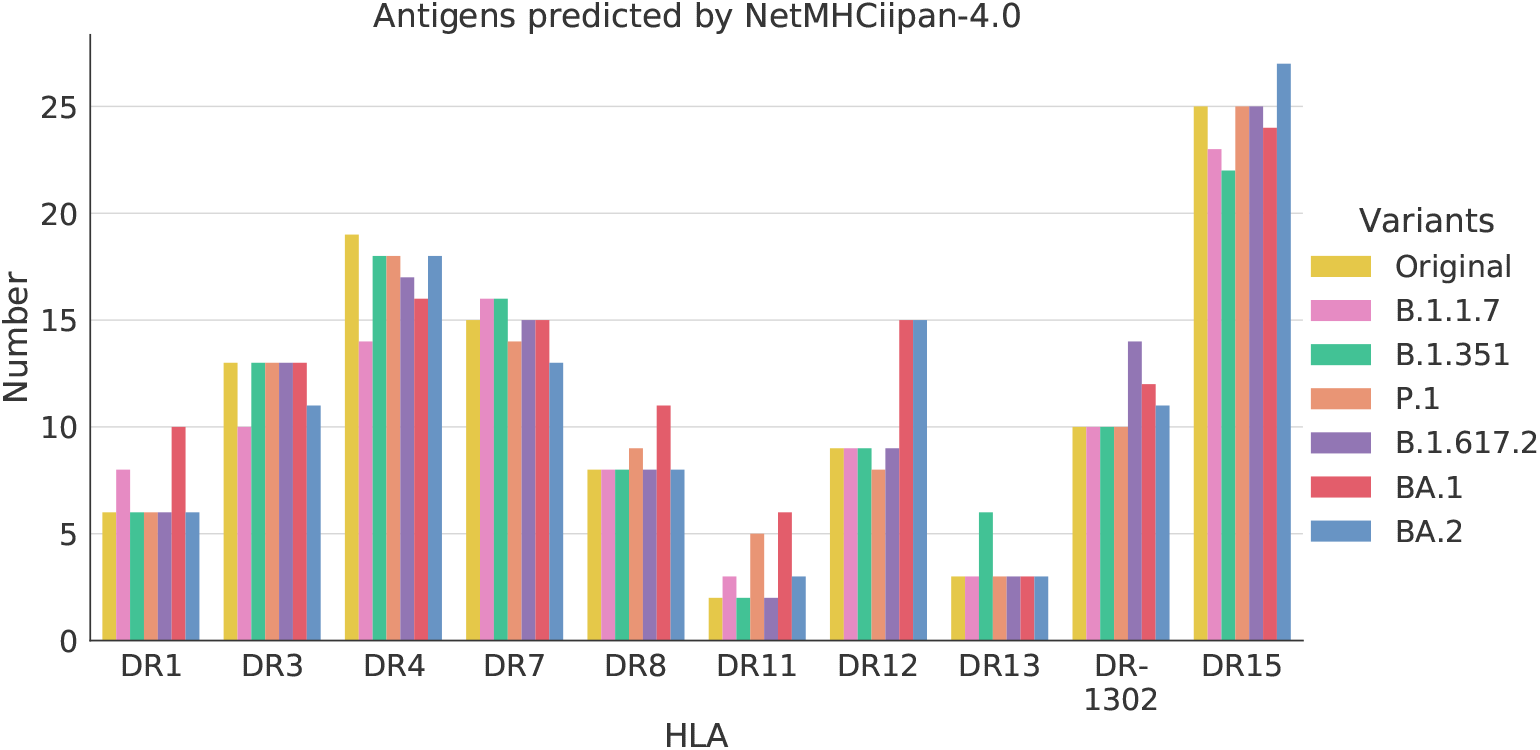
MHC Class II antigens predicted from the multiple SARS-CoV-2 variant proteins by NetMHCiipan-4.0. The x-axis tracks the various common Class II HLA types while the y-axis reports the number of antigens. The results for different variants are shown in different colored bins for each HLA.

Again, we tracked the number of antigens from the original spike that were preserved in the newer variants. These results are shown in Fig. 6. As was observed in the Class I results, the newer variants, in particular omicron, have fewer of the original antigens in several HLAs (such as DR4, DR7 and DR15). However, as the net antigenicity of each variant was relatively equivalent as seen in Fig. 5, the mutations they carried merely altered the antigens that were presented and did not lower overall antigenicity.

**Fig 6.**
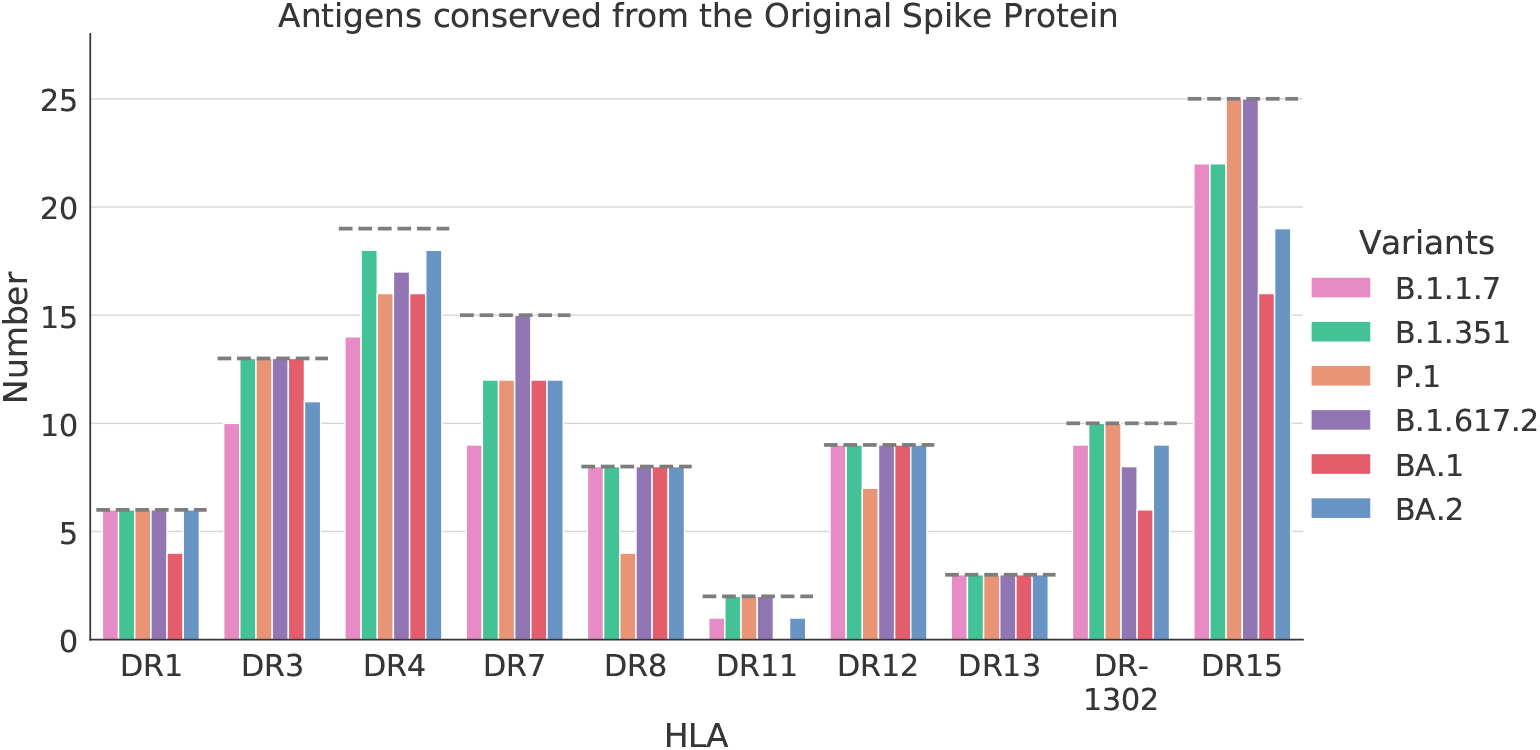
MHC Class II antigens from the original spike that were preserved in each variant spike according to NetMHCpan-4.1. For each HLA on the x-axis, the dashed line represents the number of antigens predicted from the original spike. The different variants are represented by the different colored bins. Each bin measures how many of the original spike antigens were conserved in a variant’s predicted set of antigens.

#### 3.2.2 NetMHCiipan-4.0 Results on Evaders

NetMHCiipan-4.0’s prediction results on our 100 class II evaders are shown in Fig. 7. Again, the number of predicted antigens is shown alongside the mean for all evaders and compared with the number of antigens from the original spike and the Omicron variant. For all HLAs except DR11, the sets of evaders reported a lower number of antigens than the Omicron variant. Interestingly, for DR7 and DR13 several of the evaders possessed more antigens than Omicron (for DR13, this could be because there were so few antigens already that it was easier to mutate new DR13 antigens from other HLA mutation sites in the evaders). Nonetheless, these results point out that the 36 mutations applied in most evaders led to a drop in antigenicity across all HLAs. Some HLAs, such as DR1, DR11, and DR13 even reported a complete knockout of antigens in many evaders.

**Fig 7.**
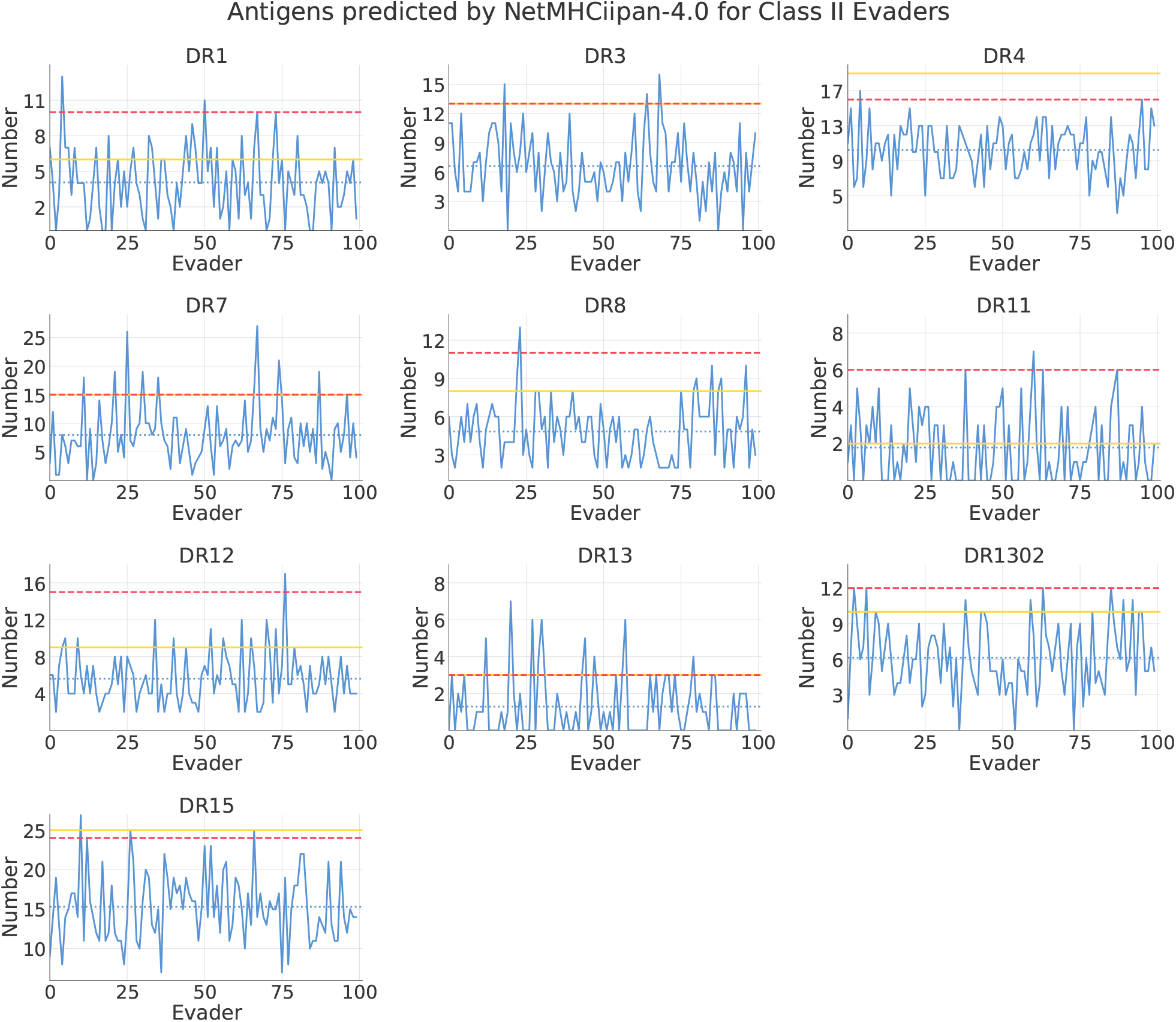
The number of strong binding peptides from each Class II evader protein predicted by NetMHCiipan-4.0 for all target HLAs. The x-axis represents all 100 evaders and the y-axis represents the number of antigens. The evader predictions are shown in the blue line plot, with the mean across all evaders in dotted blue. The number of predicted binders from the original spike is shown in solid yellow, and the number from Omicron BA.1 is shown in dashed red.

#### 3.2.3 Statistical Significance

The Wilcoxon rank-sum test results for our single class II HLA tests are shown in Table 5. Similarly to the class I results, we confirmed that the set of class II evader proteins and the set of nonrandom proteins formed statistically different distributions. The only outlier here was DR11 – we reasoned that the low number of antigens for DR11 by Omicron contributed to this discrepancy. For all the other HLAs, this suggests that our evaders significantly knocked down antigenicity of the spike protein. For these cases the omicron variant compared more favorably with the nonrandom set than the evader set as well.

Again, we investigated 3 different multi-HLA metrics. The distributions of these metrics for all evaders and nonrandom proteins is shown in Fig. 8. The first plot shows that the total antigens count of all the evaders were lower than all nonrandom proteins. The second plot shows that all evaders had a higher cartesian distance from the original spike protein’s antigen profile in comparison to the other nonrandom variants. Lastly, the third plot depicts that all evaders possessed a lower sum of ranks score than the nonrandom proteins. Again, all three metrics confirm the antigenicity knockdown across a catalogue of Class II MHc molecules by our evaders. We used the Wilcoxon rank-sum test on these metrics, as shown in Table 2, and achieved the same conclusion as the single HLA tests. The Class II evaders, with just 36 mutations from the original spike, were able to evade the MHC Class II pathway in numerous HLAs. Even in the case of DR11, the one outlier, our evaders were able to knockout antigens in several instances. These results suggest that the mutations reported in Omicron do not allow for Class II evasion.

**Fig 8.**
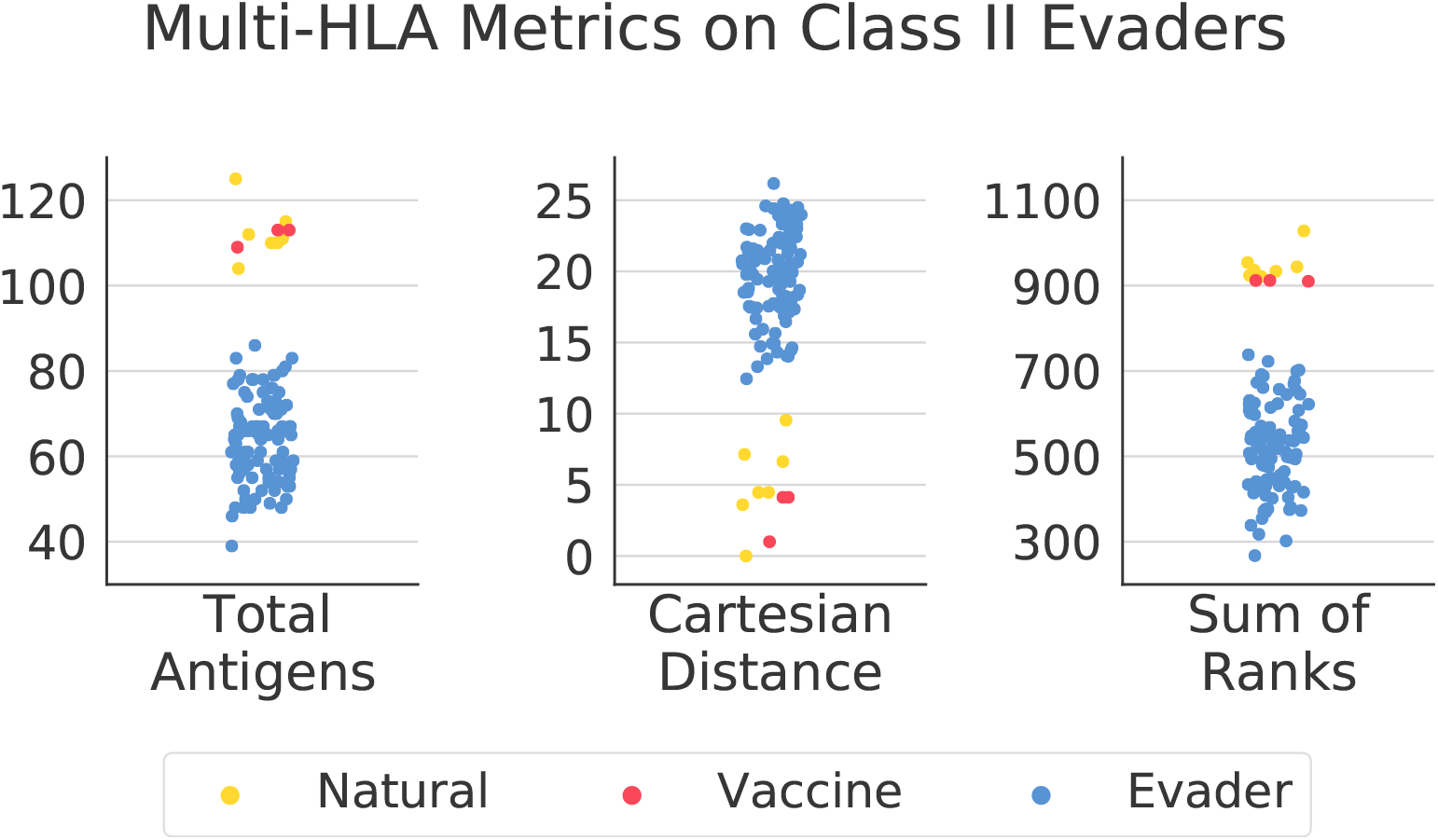
The distribution of the protein variants using the total antigens, cartesian distance, and sum of ranks metrics respectively. All Class II evaders are shown in blue, all natural SARS-CoV-2 are shown in yellow, and all vaccine spikes are shown in red. For each metric, the distributions of the evader proteins and the nonrandom (natural plus vaccine) are notably disparate.

**Table 2.** P-values for the Wilcoxon rank-sum test for multi-HLA metrics in Class II.

| Metric | Nonrandom vs. Evasive | Omicron vs. Nonrandom | Omicron vs. Evasive |
| --- | --- | --- | --- |
| Total Antigens | $2.0 \times 10^{-7}$ | $1.5 \times 10^{-1}$ | $8.6 \times 10^{-2}$ |
| Cartesian Distance | $2.0 \times 10^{-7}$ | $1.5 \times 10^{-1}$ | $8.6 \times 10^{-2}$ |
| Sum of Ranks | $2.0 \times 10^{-7}$ | $1.5 \times 10^{-1}$ | $8.6 \times 10^{-2}$ |

## Conclusion

T Cells play a vital role in immunity to viral infections. The MHC antigen presentation pathway allows for identifying more antigens, in particular peptides from the inside of a viral protein, than antibodies allow. Furthermore, T Cell levels have also been noted to last longer than antibodies levels for COVID-19 vaccine immunization [**?**]. As previous studies and our analysis have shown, T Cell immunity is particularly resilient to the numerous variants of concern for SARS-CoV-2 [15]. In particular, the two Omicron variants (BA.1 and BA.2) generated equivalent numbers of antigens in comparison to the original Wuhan spike protein according to MHC Class I and Class II antigen prediction models. Despite many peptide antigens from the original spike being lost in the newer variants, the mutations in these variants did not decrease the number of antigens across several common HLA types.

We further corroborated these observations by deliberately knocking out T Cell antigens out of the spike protein through targeted mutations. Our engineered evasive spike protein variants, despite being limited to the same number of mutations as Omicron BA.1, successfully lowered the number of antigens across numerous highly frequent HLAs. These results suggest that the mutations in the Omicron spike variant are not selected for T Cell immunity evasion, and are probably more impactful in spike protein stability or ACE2 receptor binding affinity. This explains why despite the Omicron variant being capable of neutralizing several pharmaceutical antibodies and infecting vaccinated patients, it does not evade T Cell immunity obtained by the common COVID-19 vaccines.

## Acknowledgements

We gratefully acknowledge all data contributors, i.e., the Authors and their Originating laboratories responsible for obtaining the specimens, and their Submitting laboratories for generating the genetic sequence and metadata and sharing via the GISAID Initiative, on which this research is based.

## Supporting information

### HLAs investigated

We chose the MHC Class I HLA alleles HLA-A*0101 (A1), HLA-A*0201 (A2), HLA-A*0301 (A3), HLA-A*2402 (A24), HLA-A*2601 (A26), HLA-A*3001 (A30) HLA-B*1501 (B15),HLA-B*3501 (B35), HLA-B*4001 (B40), HLA-B*4402 (B44), and HLA-B*5101 (B51).

We chose the MHC Class II HLA alleles HLA-DRB1*0101 (DR1), HLA-DRB1*0301 (DR3), HLA-DRB1*0401 (DR4), HLA-DRB1*0701 (DR7), HLA-DRB1*0801 (DR8), HLA-DRB1*1101 (DR12), DRB1*1201 (DR12), HLA-DRB1*1301 (DR13), HLA-DRB1*1302 (DR1302), and HLA-DRB1*1501 (DR15).

**Table 3.** Anchor Residue positions identified from strong binding peptides for all investigated HLAs. The amino acids that were restricted from these positions (when creating evaders) are also listed.

| HLA | Anchor Residues | Restricted Amino Acids |
| --- | --- | --- |
| A1 | 8 | Y |
| A2 | 1, 8 | L, M, I, V |
| A3 | 8 | K, R |
| A24 | 1 | Y, F, W, L |
| A26 | 8 | Y, F, M |
| A30 | 0, 2, 8 | R, K, V |
| B15 | 8 | Y, F |
| B35 | 1, 8 | P, A, Y, F, M |
| B40 | 1 | E |
| B44 | 1 | E |
| B51 | 1 | P, A |
| DR1 | 0, 3, 5, 8 | F, Y, L, G, A, V, I |
| DR3 | 3 | D |
| DR4 | 0 | F, Y, I, L |
| DR7 | 0, 3 | F, Y, I, L, S, T, V |
| DR8 | 5, 8 | K, R, D, E |
| DR11 | 5 | K, R |
| DR12 | 0, 3 | I, L, V |
| DR13 | 5 | R, K |
| DR1302 | 0, 3 | I, F, L, N, D |
| DR15 | 3 | Y, F |

**Table 4.** P-values for the Wilcoxon rank-sum test for individual HLAs in Class I. The lower the p-value, the more “separated” the two sets being compared.

| HLA | Nonrandom vs. Evasive | Omicron vs. Nonrandom | Omicron vs. Evasive |
| --- | --- | --- | --- |
| A1 | $4.3 \times 10^{-7}$ | $8.7 \times 10^{-1}$ | $9.2 \times 10^{-2}$ |
| A2 | $2.9 \times 10^{-5}$ | $8.7 \times 10^{-1}$ | $1.7 \times 10^{-1}$ |
| A3 | $2.1 \times 10^{-7}$ | $1.5 \times 10^{-1}$ | $8.6 \times 10^{-2}$ |
| A24 | $1.9 \times 10^{-9}$ | $1.5 \times 10^{-1}$ | $9.2 \times 10^{-2}$ |
| A26 | $7.0 \times 10^{-7}$ | $1.5 \times 10^{-1}$ | $1.8 \times 10^{-1}$ |
| A30 | $7.6 \times 10^{-6}$ | $1.5 \times 10^{-1}$ | $8.9 \times 10^{-2}$ |
| B15 | $3.4 \times 10^{-7}$ | $1.5 \times 10^{-1}$ | $1.7 \times 10^{-1}$ |
| B35 | $2.0 \times 10^{-7}$ | $2.6 \times 10^{-1}$ | $8.6 \times 10^{-2}$ |
| B40 | $2.7 \times 10^{-6}$ | $2.0 \times 10^{-1}$ | $1.7 \times 10^{-1}$ |
| B44 | $1.7 \times 10^{-6}$ | $1.5 \times 10^{-1}$ | $8.6 \times 10^{-2}$ |
| B51 | $1.2 \times 10^{-4}$ | $6.3 \times 10^{-1}$ | $2.4 \times 10^{-1}$ |

### New Omicron Subvariants

We used the following mutations to define the new Omicron subvariants:

1. BA.2.75: T19I, L24S, P25del, P26del, A27del, G142D, K147E, W152R, F157L, I210V, V213G, G257S, G339H, S371F, S373P, S375F, T376A, D405N, R408S, K417N, N440K, G446S, S477N, T478K, E484A, Q498R, N501Y, Y505H, D614G, H655Y, N679K, P681H, N764K, D796Y, Q954H, and N969K. There were 3 deletions and 33 point mutations.
2. BA.5: T19I, L24S, P25del, P26del, A27del, H69del, V70del, G142D, V213G, G339D, S371F, S373P, S375F, T376A, D405N, R408S, K417N, N440K, L452R, S477N, T478K, E484A, F486V, Q498R, N501Y, Y505H, D614G, H655Y, N679K, P681H, N764K, D796Y, Q954H, and N969K. There were 5 deletions and 29 point mutations.

**Table 5.** P-values for the Wilcoxon rank-sum test for individual HLAs in Class II.

| HLA | Nonrandom vs. Evasive | Omicron vs. Nonrandom | Omicron vs. Evasive |
| --- | --- | --- | --- |
| DR1 | $2.9 \times 10^{-3}$ | $1.5 \times 10^{-1}$ | $1.0 \times 10^{-1}$ |
| DR3 | $3.4 \times 10^{-6}$ | $7.5 \times 10^{-1}$ | $1.0 \times 10^{-1}$ |
| DR4 | $3.6 \times 10^{-7}$ | $2.6 \times 10^{-1}$ | $9.6 \times 10^{-2}$ |
| DR7 | $3.1 \times 10^{-5}$ | $1.0 \times 10^0$ | $1.6 \times 10^{-1}$ |
| DR8 | $1.3 \times 10^{-5}$ | $1.5 \times 10^{-1}$ | $9.2 \times 10^{-2}$ |
| DR11 | $1.1 \times 10^{-1}$ | $1.5 \times 10^{-1}$ | $1.0 \times 10^{-1}$ |
| DR12 | $5.3 \times 10^{-5}$ | $2.0 \times 10^{-1}$ | $9.2 \times 10^{-2}$ |
| DR13 | $1.9 \times 10^{-4}$ | $8.7 \times 10^{-1}$ | $2.3 \times 10^{-1}$ |
| DR1302 | $1.5 \times 10^{-5}$ | $2.6 \times 10^{-1}$ | $9.9 \times 10^{-2}$ |
| DR15 | $9.4 \times 10^{-7}$ | $4.2 \times 10^{-1}$ | $1.1 \times 10^{-1}$ |

All of these mutations were observed in more than 33% of all reported samples for that subvariant.

**Table 6.** Antigens predicted by NetMHCpan-4.1 for the vaccine and new Omicron subvariant spikes.

| HLA | A1 | A2 | A3 | A24 | A26 | A30 | B15 | B35 | B40 | B44 | B51 |
| --- | --- | --- | --- | --- | --- | --- | --- | --- | --- | --- | --- |
| BNT162b2 | 14 | 16 | 18 | 17 | 22 | 18 | 19 | 25 | 8 | 7 | 13 |
| Ad26.COV2.S | 14 | 16 | 18 | 17 | 22 | 17 | 19 | 25 | 8 | 7 | 13 |
| NVX-CoV2373 | 14 | 16 | 18 | 17 | 22 | 17 | 19 | 25 | 8 | 7 | 13 |
| BA.2.75 | 15 | 17 | 17 | 17 | 22 | 18 | 19 | 27 | 8 | 7 | 11 |
| BA.5 | 15 | 16 | 18 | 18 | 22 | 18 | 19 | 26 | 7 | 7 | 11 |

**Table 7.** Class I Antigens conserved from the Original Spike Protein in the vaccine and new Omicron subvariant spikes.

| HLA | A1 | A2 | A3 | A24 | A26 | A30 | B15 | B35 | B40 | B44 | B51 |
| --- | --- | --- | --- | --- | --- | --- | --- | --- | --- | --- | --- |
| BNT162b2 | 14 | 16 | 18 | 17 | 22 | 18 | 19 | 25 | 8 | 7 | 12 |
| Ad26.COV2.S | 14 | 16 | 18 | 17 | 22 | 17 | 19 | 25 | 8 | 7 | 12 |
| NVX-CoV2373 | 14 | 16 | 18 | 17 | 22 | 17 | 19 | 25 | 8 | 7 | 12 |
| BA.2.75 | 11 | 16 | 16 | 16 | 19 | 15 | 13 | 23 | 7 | 6 | 11 |
| BA.5 | 12 | 16 | 16 | 16 | 19 | 16 | 14 | 23 | 7 | 6 | 11 |

**Table 8.** Antigens predicted by NetMHCiipan-4.0 for the vaccine and new Omicron subvariant spikes.

| HLA | DR1 | DR3 | DR4 | DR7 | DR8 | DR11 | DR12 | DR13 | DR1302 | DR15 |
| --- | --- | --- | --- | --- | --- | --- | --- | --- | --- | --- |
| BNT162b2 | 6 | 13 | 19 | 15 | 8 | 1 | 9 | 3 | 10 | 25 |
| Ad26.COV2.S | 6 | 13 | 23 | 15 | 8 | 1 | 9 | 3 | 10 | 25 |
| NVX-CoV2373 | 6 | 13 | 23 | 15 | 8 | 1 | 9 | 3 | 10 | 25 |
| BA.2.75 | 6 | 11 | 18 | 13 | 8 | 3 | 15 | 3 | 12 | 27 |
| BA.5 | 6 | 11 | 18 | 13 | 8 | 3 | 15 | 3 | 15 | 24 |

**Table 9.** Class II Antigens conserved from the Original Spike Protein in the vaccine and new Omicron subvariant spikes.

| HLA | DR1 | DR3 | DR4 | DR7 | DR8 | DR11 | DR12 | DR13 | DR1302 | DR15 |
| --- | --- | --- | --- | --- | --- | --- | --- | --- | --- | --- |
| BNT162b2 | 6 | 13 | 19 | 15 | 8 | 1 | 9 | 3 | 10 | 25 |
| Ad26.COV2.S | 6 | 13 | 19 | 15 | 8 | 1 | 9 | 3 | 10 | 25 |
| NVX-CoV2373 | 6 | 13 | 19 | 15 | 8 | 1 | 9 | 3 | 10 | 25 |
| BA.2.75 | 6 | 11 | 18 | 12 | 8 | 1 | 9 | 3 | 5 | 19 |
| BA.5 | 6 | 11 | 18 | 12 | 8 | 1 | 9 | 3 | 7 | 16 |

## References

1. Organization WH, et al. COVID-19 weekly epidemiological update, edition 110, 21 September 2022. 2022;.

2. Khare S, Gurry C, Freitas L, Schultz MB, Bach G, Diallo A, et al. GISAID’s role in pandemic response. China CDC Weekly. 2021;3(49):1049.

3. Fan Y, Li X, Zhang L, Wan S, Zhang L, Zhou F. SARS-CoV-2 Omicron variant: recent progress and future perspectives. Signal Transduction and Targeted Therapy. 2022;7(1):1–11.

4. Sahin U, Muik A, Derhovanessian E, Vogler I, Kranz LM, Vormehr M, et al. COVID-19 vaccine BNT162b1 elicits human antibody and TH1 T cell responses. Nature. 2020;586(7830):594–599.

5. GeurtsvanKessel CH, Geers D, Schmitz KS, Mykytyn AZ, Lamers MM, Bogers S, et al. Divergent SARS-CoV-2 Omicron–reactive T and B cell responses in COVID-19 vaccine recipients. Science immunology. 2022;7(69):eabo2202.

6. Ndwandwe D, Wiysonge CS. COVID-19 vaccines. Current opinion in immunology. 2021;71:111–116.

7. Kashte S, Gulbake A, El-Amin III SF, Gupta A. COVID-19 vaccines: rapid development, implications, challenges and future prospects. Human cell. 2021;34(3):711–733.

8. Neefjes J, Jongsma ML, Paul P, Bakke O. Towards a systems understanding of MHC class I and MHC class II antigen presentation. Nature reviews immunology. 2011;11(12):823–836.

9. Bassani-Sternberg M, Chong C, Guillaume P, Solleder M, Pak H, Gannon PO, et al. Deciphering HLA-I motifs across HLA peptidomes improves neo-antigen predictions and identifies allostery regulating HLA specificity. PLoS computational biology. 2017;13(8):e1005725.

10. Gourraud PA, Khankhanian P, Cereb N, Yang SY, Feolo M, Maiers M, et al. HLA diversity in the 1000 genomes dataset. PloS one. 2014;9(7):e97282.

11. Liu L, Iketani S, Guo Y, Chan JFW, Wang M, Liu L, et al. Striking antibody evasion manifested by the Omicron variant of SARS-CoV-2. Nature. 2022;602(7898):676–681.

12. Hoffmann M, Krüger N, Schulz S, Cossmann A, Rocha C, Kempf A, et al. The Omicron variant is highly resistant against antibody-mediated neutralization: Implications for control of the COVID-19 pandemic. Cell. 2022;185(3):447–456.

13. Tao K, Tzou PL, Kosakovsky Pond SL, Ioannidis JP, Shafer RW. Susceptibility of SARS-CoV-2 Omicron variants to therapeutic monoclonal antibodies: systematic review and meta-analysis. Microbiology Spectrum. 2022;10(4):e00926–22.

14. Cox RJ, Brokstad KA. Not just antibodies: B cells and T cells mediate immunity to COVID-19. Nature Reviews Immunology. 2020;20(10):581–582.

15. Keeton R, Tincho MB, Ngomti A, Baguma R, Benede N, Suzuki A, et al. SARS-CoV-2 spike T cell responses induced upon vaccination or infection remain robust against Omicron. MedRxiv. 2021;.

16. Choi SJ, Kim DU, Noh JY, Kim S, Park SH, Jeong HW, et al. T cell epitopes in SARS-CoV-2 proteins are substantially conserved in the Omicron variant. Cellular & molecular immunology. 2022;19(3):447–448.

17. Gao Y, Cai C, Grifoni A, Müller TR, Niessl J, Olofsson A, et al. Ancestral SARS-CoV-2-specific T cells cross-recognize the Omicron variant. Nature medicine. 2022;28(3):472–476.

18. Naranbhai V, Nathan A, Kaseke C, Berrios C, Khatri A, Choi S, et al. T cell reactivity to the SARS-CoV-2 Omicron variant is preserved in most but not all individuals. Cell. 2022;185(6):1041–1051.

19. Tarke A, Sidney J, Methot N, Yu E, Zhang Y, Dan J, et al.. Impact of SARS-CoV-2 variants on the total CD4 (+) and CD8 (+) T cell reactivity in infected or vaccinated individuals. Cell Rep Med. 2021; 2 (7): 100355; 2021.

20. Reynisson B, Alvarez B, Paul S, Peters B, Nielsen M. NetMHCpan-4.1 and NetMHCIIpan-4.0: improved predictions of MHC antigen presentation by concurrent motif deconvolution and integration of MS MHC eluted ligand data. Nucleic acids research. 2020;48(W1):W449–W454.

21. Paul S, Grifoni A, Peters B, Sette A. Major histocompatibility complex binding, eluted ligands, and immunogenicity: benchmark testing and predictions. Frontiers in immunology. 2020;10:3151.

22. McGranahan N, Rosenthal R, Hiley CT, Rowan AJ, Watkins TB, Wilson GA, et al. Allele-specific HLA loss and immune escape in lung cancer evolution. Cell. 2017;171(6):1259–1271.

23. Łuksza M, Riaz N, Makarov V, Balachandran VP, Hellmann MD, Solovyov A, et al. A neoantigen fitness model predicts tumour response to checkpoint blockade immunotherapy. Nature. 2017;551(7681):517–520.

24. Zhou P, Yang XL, Wang XG, Hu B, Zhang L, Zhang W, et al. A pneumonia outbreak associated with a new coronavirus of probable bat origin. nature. 2020;579(7798):270–273.

25. Okada P, Phuygun S, Thanadachakul T, Parnmen S, Wongboot W, Waicharoen S, et al. Early transmission patterns of coronavirus disease 2019 (COVID-19) in travellers from Wuhan to Thailand, January 2020. Eurosurveillance. 2020;25(8):2000097.

26. Walls AC, Park YJ, Tortorici MA, Wall A, McGuire AT, Veesler D. Structure, function, and antigenicity of the SARS-CoV-2 spike glycoprotein. Cell. 2020;181(2):281–292.

27. Gangavarapu K, Latif AA, Mullen JL, Alkuzweny M, Hufbauer E, Tsueng G, et al. Outbreak.info genomic reports: scalable and dynamic surveillance of SARS-CoV-2 variants and mutations. Research square. 2022;.

28. Tsueng G, Mullen J, Alkuzweny M, Cano M, Benjamin HR, Emily O, et al. Outbreak. info Research Library: A standardized, searchable platform to discover and explore COVID-19 resources and data. BioRxiv. 2022;.

29. Hodcroft EB. CoVariants: SARS-CoV-2 mutations and variants of interest; 2021.

30. Gerdol M, Dishnica K, Giorgetti A. Emergence of a recurrent insertion in the N-terminal domain of the SARS-CoV-2 spike glycoprotein. Virus research. 2022;310:198674.

31. Maiers M, Gragert L, Klitz W. High-resolution HLA alleles and haplotypes in the United States population. Human immunology. 2007;68(9):779–788.

32. Hoof I, Peters B, Sidney J, Pedersen LE, Sette A, Lund O, et al. NetMHCpan, a method for MHC class I binding prediction beyond humans. Immunogenetics. 2009;61(1):1–13.

33. Reynisson B, Barra C, Kaabinejadian S, Hildebrand WH, Peters B, Nielsen M. Improved prediction of MHC II antigen presentation through integration and motif deconvolution of mass spectrometry MHC eluted ligand data. Journal of proteome research. 2020;19(6):2304–2315.

34. Grifoni A, Weiskopf D, Ramirez SI, Mateus J, Dan JM, Moderbacher CR, et al. Targets of T cell responses to SARS-CoV-2 coronavirus in humans with COVID-19 disease and unexposed individuals. Cell. 2020;181(7):1489–1501.

